# Dentate gyrus somatostatin cells are required for contextual discrimination during episodic memory encoding

**DOI:** 10.1101/830182

**Authors:** Cristian Morales, Juan Facundo Morici, Nelson Espinosa, Agostina Sacson, Ariel Lara-Vasquez, M. Alexandra Garcia-Perez, Pedro Bekinschtein, Noelia V. Weisstaub, Pablo Fuentealba

## Abstract

Episodic memory establishes and stores relations among the different elements of an experience, which are often similar and difficult to distinguish. Pattern separation, implemented by the dentate gyrus, is a neural mechanism that allows the discrimination of similar experiences by orthogonalizing synaptic inputs. Granule cells support such disambiguation by sparse rate coding, a process tightly controlled by highly diversified GABAergic neuronal populations, such as somatostatin-expressing cells which directly target the dendritic arbor of granule cells, massively innervated by entorhinal inputs reaching the molecular layer and conveying contextual information. Here, we tested the hypothesis that somatostatin neurons regulate the excitability of the dentate gyrus, thus controlling the efficacy of pattern separation during memory encoding in mice. Indeed, optogenetic suppression of dentate gyrus somatostatin neurons increased spiking activity in putative excitatory neurons and triggered dentate spikes. Moreover, optical inhibition of somatostatin neurons impaired both contextual and spatial discrimination of overlapping episodic-like memories during task acquisition. Importantly, effects were specific for similar environments, suggesting that pattern separation was selectively engaged when overlapping conditions ought to be distinguished. Overall, our results suggest that somatostatin cells regulate excitability in the dentate gyrus and are required for effective pattern separation during episodic memory encoding.

**Significance statement:** Memory systems must be able to discriminate stored representations of similar experiences in order to efficiently guide future decisions. This is solved by pattern separation, implemented in the dentate gyrus by granule cells to support episodic memory formation. The tonic inhibitory bombardment produced by multiple GABAergic cell populations maintains low activity levels in granule cells, permitting the process of pattern separation. Somatostatin-expressing cells are one of those interneuron populations, selectively targeting the distal dendrites of granule cells, where cortical multimodal information reaches the dentate gyrus. Hence, somatostatin cells constitute an ideal candidate to regulate pattern separation. Here, by using optogenetic stimulation in mice, we demonstrate that somatostatin cells are required for the acquisition of both contextual and spatial overlapping memories.

## Introduction

Episodic memory is made up of a collection of different events which may contain overlapping information. The capacity to discriminate similar memory episodes, which could otherwise be confused, is critical for correct encoding (1), effective retrieval (2, 3), and avoiding catastrophic interference (4). Accordingly, disruptions of memory discrimination have been related to cognitive impairment in aging and neuropsychiatric disorders, such as depression or posttraumatic stress disorder (5–7). In addition, aging and epilepsy exhibit episodic memory deficits as very common symptoms, also showing disrupted pattern separation (8–10). Computational models suggest that pattern separation ought to be implemented for efficient discrimination of overlapping memories, by the transformation of similar patterns of inputs into segregated synaptic outputs (11–13). Abundant evidence from animal experiments support the hippocampus as the brain locus where pattern separation is implemented, in particular the dentate gyrus, which allows the discrimination of overlapping spatial representations (14–16) and similar contextual fear conditioning memories (17).

Subsets of granule cells in the dentate gyrus represent memory episodes in their activation patterns (18, 19), with similar episodes of experience being stored into distinct non-overlapping activation patterns in a process called orthogonalization (20–22) that is essential for pattern separation. Granule cells conform a large neuronal population that remains sparsely active due to strong inhibitory control provided by local GABAergic inputs (23–25). The tight regulation of granule cell excitability is critical (26, 27), as evidenced by deteriorated pattern separation when the spiking activity of granule cells is experimentally enhanced (28) or when tonic inhibition is decreased. Moreover, tonic inhibition of dentate gyrus granule cells is important for the control of memory interference (29). The most significant inhibitory inputs to granule cells arise locally from parvalbumin-expressing (PV) cells and somatostatin-expressing neurons (SOM) located mostly in the hilus (28, 30, 31). PV cells provide massive synaptic inhibition to the perisomatic region (32), whereas SOM directly innervate the dendritic arbor of granule cells. Granule cell dendritic arborization is largely distributed across the molecular layer, and it is this region where entorhinal inputs massively impinge carrying contextual information (30). Recent experiments have changed the view about the modulatory function of SOM and established their control over the excitability of granule cells in vitro (33) and the size of neuronal ensembles during memory encoding in vivo (34). Taken together, these findings suggest that SOM are ideally positioned to participate in contextual discrimination, possibly through the regulation of pattern separation by controlling the dendritic excitability of granule cells (8, 35). Hence, we tested the hypothesis that functional dentate gyrus SOM are required for contextual discrimination through the control of granule cells excitability. We found that optogenetic suppression of SOM in the dentate gyrus modulated the firing rate of putative excitatory cells and putative PV interneurons. Moreover, optogenetic stimulation impaired both contextual and spatial discrimination of overlapping recognition memories during task acquisition. Our results suggest that SOM are required for successful pattern separation during episodic memory encoding.

## Results

### Optogenetic suppression of somatostatin cells disrupts dentate gyrus neuronal firing patterns and trigger dentate spikes

In order to establish the effect of locally inhibiting SOM on the dentate gyrus network, we stereotaxically implanted an optrode covering the entire dorsoventral extension of the dentate gyrus (**Fig. S1**) in anesthetized double transgenic mice that selectively expressed the inhibitory halorhodopsin pump (NpHR+) in SOM (36, 37). To ensure the simultaneous recording of all three dentate gyrus laminae and optical suppression of hilar SOM (38), the tip of the optical fibre was positioned close to the hippocampal fissure (**Fig. 1A**). Next, we delivered prolonged laser pulses to achieve maximal optogenetic inhibition, reproducing previous experimental protocols (36, 37). Control experiments showed that effects were selective for transgenic NpHR+ animals, with little effect on control (NpHR-) mice (**Fig. S1**).

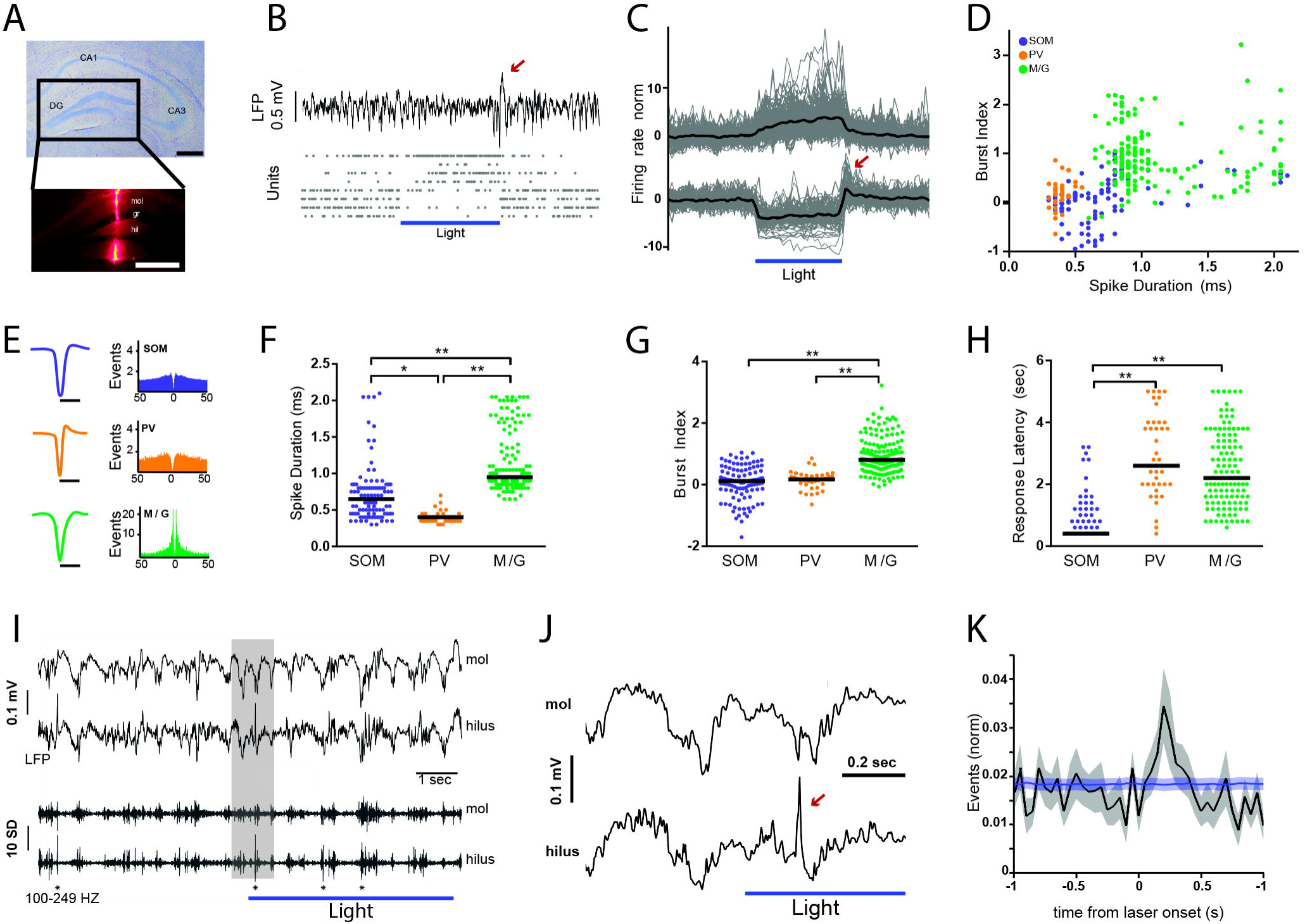

Overall, we found a small proportion of units changing their firing patterns upon laser stimulation in NpHR+ mice. A minor fraction of units (6.3%, n = 111) robustly decreased (median = 57%, IQR = 55%) their activity relatively fast (median = 400 ms, IQR = 200 ms). Hence, such neuronal population was defined as SOM (**Fig. 1C**). Another small group of cells (12 %, n = 203) showed a significant increase in firing rate (median = 87%, IQR 127%), presumably by synaptic disinhibition, with significantly slower kinetics than SOM (median = 2.4 s, IQR = 2 s, P < 0.001, **Fig. 1C, Fig. S1**). Dentate gyrus units have been previously classified as glutamatergic or GABAergic based on their in vivo spike duration and bursting patterns (39). Indeed, excitatory cells, such as mossy cells and granule cells exhibit longer trough-to-peak latency and higher burst index than GABAergic interneurons (39). Thus, we carried out a similar analysis. We first plotted all laser-responsive units in a bidimensional space comprised by spike duration and burst index (**Fig. 1D**). By definition, SOM comprised one cluster given their common physiological characteristic of optogenetic inhibition (**Fig. 1D,E**). Within the optically excited units we recognized two different populations. We considered one group of excited units as putative PV cells because of their short spikes, significantly more rapid than the other groups (median = 0.4 ms, IQR = 0.1 ms, P < 0.001; **Fig. 1F**). Such short spike duration has been well documented in GABAergic PV cells (39–41) that are synaptically targeted by SOM (42). On the other hand, the other cluster of excited units was consistent with the presence of mossy cells and granule cells (M/G). Indeed, M/G units had slower spikes (median = 0.95 ms, IQR = 0.34 ms, P < 0.001; **Fig. 1F**) and higher burst index (median = 0.8, IQR = 0.6, P < 0.001; **Fig. 1G**) than either PV cells or SOM. Furthermore, there was no significant difference in response latency to optogenetic stimulation between PV cells and M/G units (**Fig. 1H**), but they were both significantly slower than SOM (median = 0.4 s, IRQ = 0.2 s, P < 0.001; **Fig. 1H**). These results suggest that a small proportion of both PV cells and M/G units was probably disinhibited upon optogenetic suppression of SOM.

Next, we aimed to evaluate the effect of the modulated synaptic output of SOM onto dentate gyrus oscillatory activity. We noted that at the end of laser pulses a prominent deflection was apparent in the dentate gyrus field potential (**Fig. 1B**) that was probably the result of rebound activity in SOM (**Fig. 1C**). A similar effect has been previously described in the dorsal CA1 area (43). We then quantified the spectral distribution of dentate gyrus activity. The LFP frequency spectrum showed a prominent shoulder in the gamma range (30-80 Hz) which then decayed with the characteristic 1/f distribution at higher frequencies ((44), **Fig. S2**). Interestingly, optogenetic suppression of SOM selectively decreased power in the high-frequency range (100-250 Hz, **Fig. S2**). Dentate spikes are hallmark population patterns exclusive of the dentate gyrus, which due to their large-amplitude, short-duration can be detected in the high-frequency activity range (45). Indeed, high-pass filtering LFP recordings, particularly in the hilar region, evidenced the presence of prominent dentate spikes (**Fig. 1I,J**). Dentate spikes result from massive dendritic depolarization and perisomatic inhibition of granule cells (45, 46). Given that optogenetic suppression of SOM increased the activity of putative granule cells and PV interneurons, that inhibit the perisomatic region of granule cells (38), we reasoned that during optical stimulation the incidence of dentate spikes was likely to increase. Indeed, crosscorrelation analysis between laser pulses and dentate spikes showed a significant increase in their probability of occurrence (**Fig. 1K**). Similarly, their incidence increased when comparing the periods before and after the onset of laser stimulation (**Fig. S2**). Hence, the inhibition of SOM may disinhibit PV interneurons and granule cells, with the concomitant increase of network excitability, reflected in the elevated density of dentate spikes. Altogether, our results suggest that optogenetic inhibition of SOM disrupts local network activity patterns in the dentate gyrus.

### Encoding of overlapping contextual memories requires functional glutamatergic transmission and somatostatin cells in the dentate gyrus

Fully functional granule cells are necessary for the discrimination of similar contexts (17, 47) as well as the acquisition of novel information. However, the role of other dentate gyrus cell-types is not so well established, particularly interneuron populations that regulate the spike timing of granule cells. Hence, we investigated the contribution of hilar SOM to behavioural performance in context-dependent memory tasks. To test this idea, we developed a variation of the object-in-context task (48–52). We reasoned that discrimination of the novel object-context pairing should recruit the dentate gyrus only when contextual configurations were overlapping. Accordingly, we manipulated contextual information by training animals in the object-in-context task using similar or dissimilar contexts (**Fig. 2A, Fig. S3**). First, we evaluated whether the dentate gyrus was engaged in contextual memory encoding in this test. For this, we blocked glutamatergic transmission during memory encoding by infusing DNQX into the dentate gyrus preceding every training session. Regardless of the object-context configuration, blockade of AMPA receptors did not affect exploratory behaviour during training sessions (**Fig. S4**). Consistent with our hypothesis, blocking dentate gyrus excitatory transmission obliterated discrimination during the test only when contexts were similar (mean = 0.013, SEM = 0.012, P < 0.001; **Fig. 2C,E, Fig. S5**). Importantly, the experimental manipulation did not affect total exploration time (**Fig. 2D,F**), suggesting that these results were not due to changes in motivation, awareness, or exploratory behaviour in general. Together, these results suggest that glutamatergic transmission in the dentate gyrus is necessary for the encoding of object-in-context memories only when the contextual information presented is similar, consistent with the observation that pattern separation is engaged only when overlapping spatial representations ought to be discriminated (14, 53).

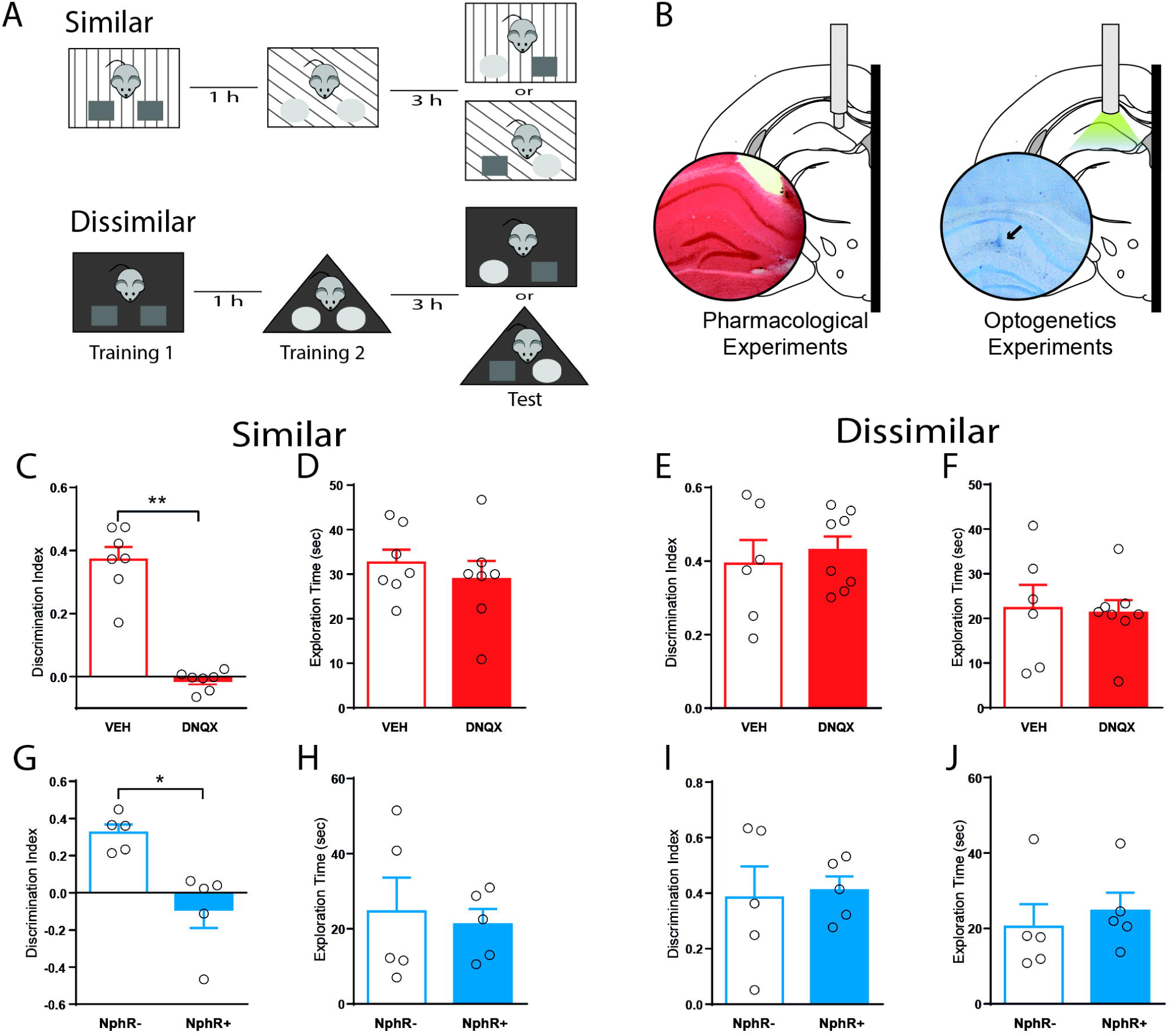

It has also been proposed that the dentate gyrus is recruited during the retrieval of similar representations (54–56). Therefore, we manipulated glutamatergic transmission in the dentate gyrus during memory recall (**Fig. S6**). For that purpose, we performed the object-in-context task in the similar condition and infused DNQX into the dentate gyrus 15 minutes before the test session, when memory is retrieved. In this condition, we did not detect any effect of the drug on discrimination. Interestingly, blocking AMPA receptors influenced behaviour as exploration times were higher for drug-infused animals (mean = 33.83 s, SEM = 1.95 s, P < 0.001; **Fig. S6**). Hence, this result suggests that the dentate gyrus is actively engaged during encoding of similar contextual representations, but not during their retrieval.

Recent evidence suggests that SOM are recruited during fear memory formation (34, 57) and may participate in pattern separation by controlling the size of engrams encoding spatial representations (34, 38, 58). Consequently, we conducted the object-in-context task in our transgenic mice expressing functional halorhodopsin (NpHR+) in SOM (36). We bilaterally implanted optic fibres into the dentate gyrus (**Fig. 2B**) in order to activate NpHR during the training sessions (**Fig. 2A**). Laser stimulation did not affect the distribution of object exploration times during training sessions for neither task version (**Fig. 2H,J, Fig. S4**). Nonetheless, optogenetic inhibition of SOM during the training session impaired discrimination during retrieval when contexts were similar (mean = 0.09, SEM = 0.098, P < 0.01; **Fig. 2G, Fig. S5**). This effect was specific, as it was not detected for the dissimilar context (**Fig. 2I, Fig. S5**), suggesting that dentate gyrus SOM regulate the encoding of object-in-context memory only when contextual information is similar. Overall, this result suggests that SOM regulate the encoding of contextual recognition memory by controlling excitatory activity in the dentate gyrus.

### Discrimination of overlapping spatial configurations is regulated by somatostatin neurons during memory encoding

We showed that SOM can control the encoding of contextual recognition memory. Previous studies have shown the essential role played by granule cells in encoding distinct neural representations of overlapping spatial configurations (20, 22, 59). Furthermore, inhibition of the lateral entorhinal input disrupts pattern separation in spatial tasks (60). Therefore, we reasoned that hilar SOM may also regulate the encoding of overlapping spatial configurations. To test this, we conducted the spontaneous location recognition task that has been shown to be sensitive to the functional integrity of the dentate gyrus (16). In this task animals explore three identical objects placed in separate locations and are then tested with two objects, with one object placed in a familiar location and another object placed in between the previous two locations (**Fig. 3A**). As the two objects are placed closer together in the training session, it becomes more difficult for mice to discriminate the novel location in the test session (16, 61, 62). Accordingly, we varied the angle between objects during the training sessions in small (50 degrees), medium (120 degrees), or large (180 degrees) separations; and compared the exploration times between the training and test session. We found that mice were able to discriminate correctly the medium and large separations, yet failed to distinguish the small separation (mean = -0.062, SEM = 0.068, P < 0.05; **Fig. 3B**). This effect was robustly expressed during the entire test session (**Fig. S7**). These results suggest that the small separation is too ambiguous for mice to be able to discriminate it. Thus, we assessed whether spatial discrimination of the medium configuration relied on the activity of hilar SOM. For that, we chronically implanted bilateral optic fibres on the hippocampal fissure of transgenic NpHR+ mice (**Fig. 3D**) to optogenetically inhibit SOM during memory encoding. Laser stimulation had no effect on exploratory behaviour during the training sessions (**Fig. S7**). However, optogenetic inhibition of SOM during acquisition selectively impaired spatial discrimination in NpHR+ mice without affecting performance in control NpHR-mice (mean = 0.014, SEM = 0.012, P < 0.05; **Fig. 3E**). Importantly, exploratory behaviour of mice was not affected by laser stimulation (**Fig. 3F**). This result suggests that SOM regulate the encoding of spatial recognition memory by controlling neural activity in the dentate gyrus. Overall, our results suggest that SOM can regulate both the reactivity and excitability of granule cells in-vivo (**Fig. S8**).

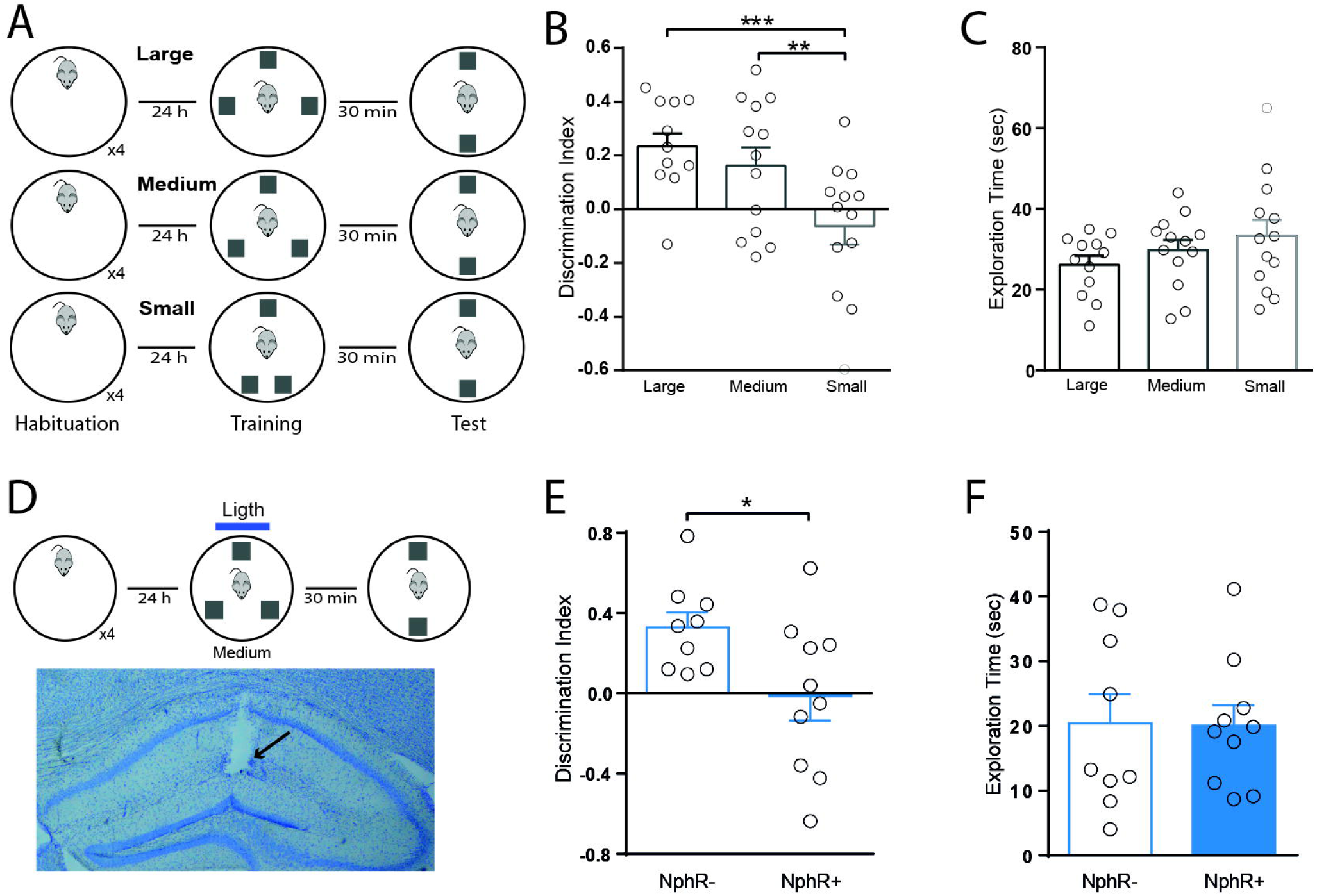

## Discussion

Here, we studied the physiological and behavioral effects of selectively suppressing dentate gyrus SOM during contextual memory acquisition. We found that inhibition of SOM increased the activity of two neuronal populations identified on the basis of their brief spike waveforms and firing patterns. Indeed, inhibitory interneurons, putative PV cells, were identified by their fast spiking pattern and short spikes (39–41); whereas, excitatory neurons, putative granule cells and mossy cells, were recognised by their bursting pattern and slow spikes (39). Both neuronal populations increased their activity during laser stimulation, probably resulting from synaptic disinhibition (30), which was also consistent with the enhanced incidence of dentate spikes, a hallmark of dentate gyrus synchronous activity (45, 46). Low excitability of granule cells has been proposed as a requirement for efficient orthogonalization of afferent spiking patterns arising in the entorhinal cortex (27). Thus, nonselective increments of activity in granule cells are predicted to alter efficient pattern separation. By performing two types of episodic-like memory tasks, one contextual and another spatial, we reveal that discrimination of similar memories is disrupted by optogenetic inhibition of dentate gyrus SOM.

The hippocampus has been proposed as a key structure for the control of memory expression using contextual information (63). For this reason, the ability to differentiate overlapping contextual information during encoding should be important to facilitate posterior memory recall (64). In this regard, previous studies have typically used the fear conditioning paradigm to characterize the role of the dentate gyrus in encoding overlapping contextual information (17, 65). Given the emotional valence of this kind of task, the amygdala is likely to be recruited (66), and possibly activate the hippocampus. Indeed, the basolateral amygdala is densely interconnected with the ventral hippocampus (67–69). Thus, engagement of the dentate gyrus is expected when the amygdala is massively activated during aversive memory formation. The dentate gyrus also participates in the differentiation of overlapping object-spatial representations in neutral conditions (16, 62). By experimentally manipulating object-context associations we establish here that excitatory activity in the dentate gyrus mediated by AMPA receptors is necessary to discriminate overlapping contextual associations. Our results suggest that the dentate gyrus is involved in the general process of discriminating overlapping spatial-contextual information independently of its emotional valence. Specifically, we show that inhibition of hilar SOM impaired the acquisition of similar contextual and spatial representations, thus suggesting that somatostatin-dependent inhibition of the dentate gyrus is at play in the discrimination of different types of similar episodic memories. During pattern separation, different neuronal engrams represent similar contexts (59). It has been shown that some of the dentate gyrus cells are highly sensitive to small changes in contextual cues (20, 70), suggesting that dentate gyrus ensembles recognize differences in contextual information with high sensitivity. Importantly, SOM regulate the size of such memory ensembles (34), and thus inhibiting SOM is expected to dysregulate both the size and specificity of memory engrams. This hypothesis is supported by the anatomical distribution of SOM that selectively target the most external region of the dendritic trees of granule cells in the outer molecular layer (38), where contextual information is conveyed by entorhinal inputs (58), and damage to those afferents impairs spatial pattern separation (60). Thus, the behavioral impairment resulting from the optogenetic inhibition of SOM could be attributed to deficits in the ability of the dentate gyrus to differentially encode contextual information resulting from impaired pattern separation.

Pattern separation is believed to be critical during contextual memory acquisition, but its contribution to memory recall remains debated. It has been proposed that inhibition and excitation of dentate gyrus circuits play different roles during encoding, consolidation, or recall of overlapping memories (71, 72). This is supported by the blockade of dentate gyrus AMPA and kainite receptors, which impairs the expression of fear memory (73), while the blockade of GABA_A_ receptors impairs consolidation, yet not the acquisition or retrieval of fear memories (74). Some theoretical models propose that sparse patterns of neuronal activation in the dentate gyrus guide memory encoding in the CA3 region, whereas memory retrieval is mediated through direct entorhinal inputs to the CA3 region (26). However, other studies suggest that the dentate gyrus contributes to both memory encoding and retrieval (54–56). This is supported by recent studies showing that recall cues can trigger reactivation of neural ensembles active in the dentate gyrus during memory encoding (19, 75, 76). Interestingly, we found that blockade of dentate gyrus AMPA receptors did not affect the recall of contextual memories. The differences observed could be, at least partly, due to methodological reasons. For example, we pharmacologically blocked AMPA receptors before the test session whereas the above-mentioned studies selectively controlled very specific subsets of dentate gyrus neurons. Hence, our results support the idea that the role of the dentate gyrus in the process of pattern separation is restricted to the encoding phase, at least, for emotionally-neutral memories.

Feedback inhibition in the dentate gyrus is necessary for appropriate orthogonalization of neuronal ensembles representing similar memory episodes (77). SOM receive direct inputs from both mature (38) and newborn (78) granule cells. Both granule cell populations have been proposed as drivers of feedback inhibition in pattern separation (34, 79). Several studies proposed a leading role for newborn granule cells (77, 79). Indeed, the activation of newborn granule cells engages inhibitory feedback principally arising from PV interneurons that provide perisomatic inhibition, but not from SOM that provide dendritic inhibition (80). Moreover, ablation of hippocampal neurogenesis induces high excitability in granule cells (81) and impairs pattern separation (15), supporting the idea that feedback inhibition mediated by PV interneurons controls pattern separation. On the other hand, a recent study suggests that mature granule cells poorly engage PV interneurons and preferentially excite SOM in vivo (34). Thus, it is plausible that in pattern separation two parallel circuits are engaged during feedback inhibition. One circuit in which newborn granule cells preferentially recruit PV cells and another circuit where mature granule cells preferentially activate SOM that would control the inhibition of granule cells directly. Since newborn cells are more excitable that granule cells (82), it is possible that feedback activity of SOM is more delayed than feedback inhibition of PV cells. Interestingly, computational models of pattern separation predict little contribution for the inhibitory feedback provided by SOM (77). Our results suggest that feedback inhibition provided by SOM is necessary for pattern separation and it will be interesting that future computational studies consider this parameter when modelling hippocampal networks.

Furthermore, the regulation of orthogonalization of granule cells relies heavily on the perisomatic lateral inhibition provided by parvalbumin cells (83, 84). In turn, lateral inhibition might be controlled by SOM through direct dendritic inhibition of granule cells and perisomatic inhibition of parvalbumin cells (30). When mice explore novel environments, thus forming new memories, or during intense presynaptic activity in the dentate gyrus in vitro, dendritic inhibition is significantly larger than perisomatic inhibition (85, 86). A population of SOM locally innervates fast-spiking PV cells in the dentate gyrus and distally several other cell-types in the septum (30). Consistent with such anatomic connectivity, we found that inhibition of SOM increased the activity of putative PV neurons and granule cells. Moreover, we observed increased incidence of dentate spikes, which results from simultaneous, brief dendritic depolarization and perisomatic inhibition of granule cells (45, 46). Furthermore, SOM synaptically target PV interneurons unidirectionally, with no feedback projection described to date (42, 87). This synaptic projection regulates both the discharge probability and spike timing of PV neurons (30, 42). These changes in the dentate gyrus circuit are consistent with memory deficits and disrupted pattern separation taking place in cases where the population of hilar SOM is selectively decreased, such as epilepsy or aging (6, 10, 88). In summary, our results suggest that SOM regulate general excitability in the dentate gyrus and are required for pattern separation during episodic memory encoding.

## Material and methods

95 adult male and female (C57Bl/6J, Ai39, B6, 129S-Gt(ROSA)26Sor^tm39(CAG-HOP/EYFP)Hze^/J, Sst-IRES-Cre, Sst^tm2.1(cre)Zjh^/J and B6N.Cg-Sst^tm2.1(cre)Zjh^/J) were used. All transgenic lines were obtained from Jackson laboratories (www.jax.org). Mice were deeply anesthetized with urethane and acutely implanted with multielectrode probes (www.neuronexus.com) targeting the dorsal dentate gyrus (−2.0 mm AP, 1.5 mm ML from Bregma) to record neural activity while SOM were optogenetically suppressed with laser stimulation (532 nm, 5-12 mW). Particularly, 33 mice were chronically implanted with optic fibres or cannulated in the dentate gyrus and trained in contextual and spatial object recognition memory tasks. In both tasks, mice were first habituated to the arena, next trained in the task, and then tested in the final session. In some sessions, animals were locally injected in the dentate gyrus with an AMPA receptor blocker (DNQX, 1.89 μg/μl, 0.3 ul). Behavioral analysis included quantification of exploration time and discrimination index. Spectral analysis of LFP was computed using multitaper Fourier analysis from the Chronux toolbox (http://www.chronux.org) by using MATLAB. Spike sorting and single unit analysis was performed offline using MATLAB based graphical cluster-cutting software, Mclust/Klustakwik-toolbox (version 3.5; (89)). To verify recording sites, optic fibers and cannulae tracks, Nissl-staining was conducted following the completion of study. See Supplementary Materials and Methods section for a detailed description of the experimental and analytic methods.

## Acknowledgments

We thank Maria José Díaz and David Alberto Jaime for their technical assistance.

