## supplementary figures for "Dentate gyrus somatostatin cells are required for contextual discrimination during episodic memory encoding"

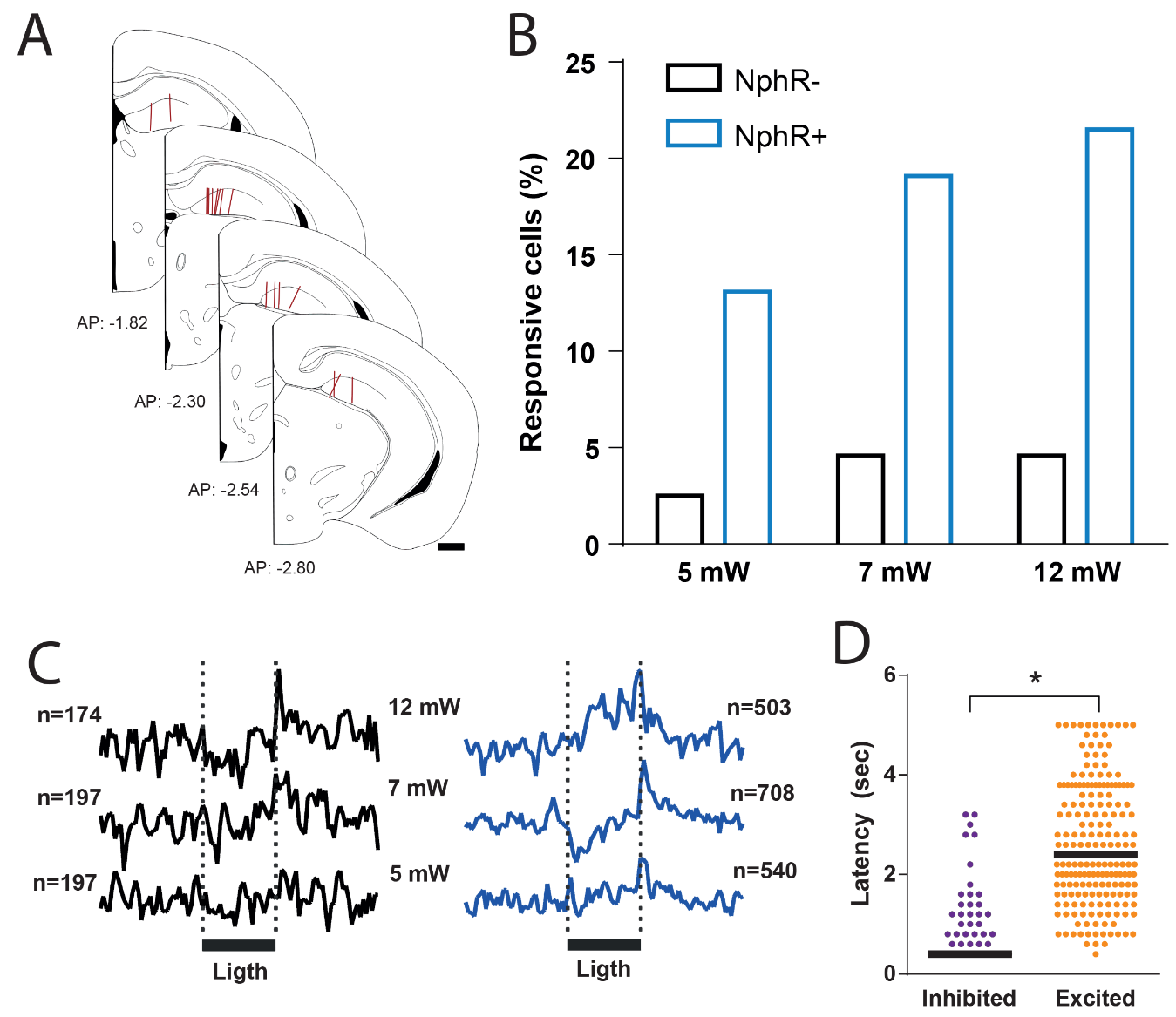

**Supplementary Figure 1. Neuronal spiking responses to optogenetic stimulation in the hippocampus. A,** Anatomical tracks (red lines) of optrodes positioned in the dentate gyrus. Scale bar, 500 µm. **B,** Proportion of units responsive to optogenetic stimulation at different power intensities in transgenic mice expressing halorhodopsin (NphR+) or control mice (NphR-). Proportions were significantly different between NphR+ or NphR- mice. X^2^ test; 5 mW, P < 10^-4^; 7 mW, P < 10^-6^; 12 mW, P < 10^-6^. **C,** Normalized average discharge probability for all recorded units in NphR- mice (black lines) and NphR+ mice (blue line). Black horizontal bar depicts laser stimulation (5 seconds, fiber diameter 100 um). **D,** Response latency of hippocampus units to laser stimulation. Latency was significantly larger in excited units than inhibited units (Wilcoxon rank sum test, * P = 7.8x10^-39^), thus suggesting different synaptic connectivity.

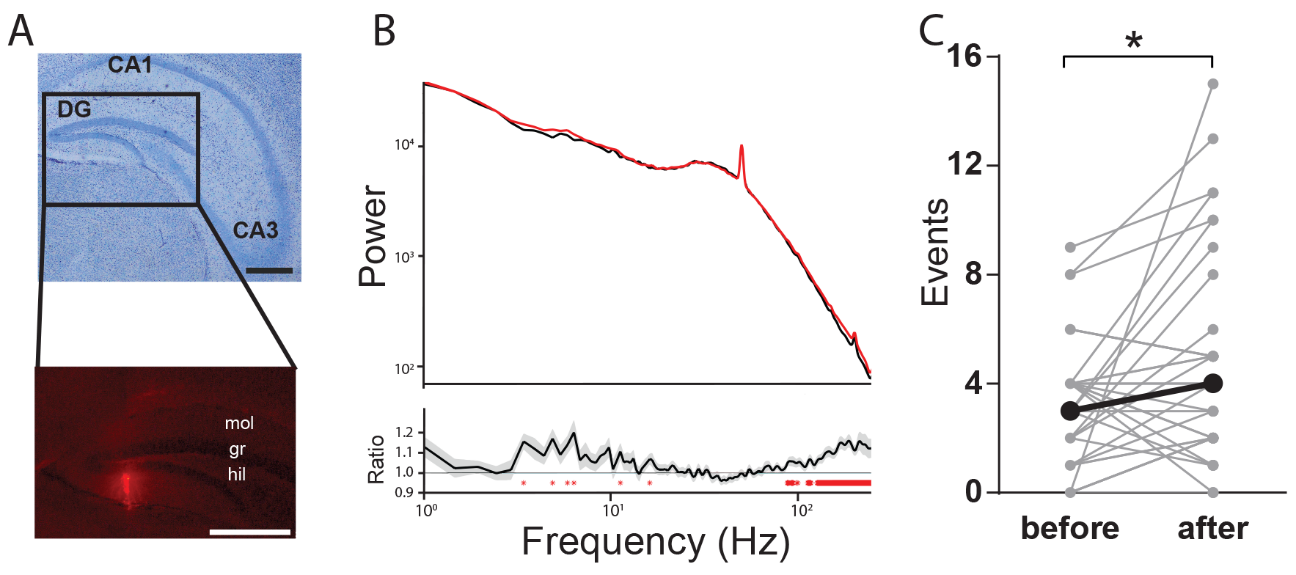

**Supplementary Figure 2. Field potential changes in response to optogenetic stimulation in the hippocampus. A,** Nissl-stained brain section of dorsal hippocampus showing different regions, CA1, CA3 and dentate gyrus (DG). Scale bar, 500 µm. Inset: fluorescence micrography of the DG with optrode track stained with DiI. mol, molecular layer; gr., granular layer; hil., hilus. Scale bar, 500 µm. **B,** Power spectral distribution of granular layer activity during light-on periods (red line) contrasted with light-off periods (black line). Bottom, ratio between light-on periods (red line) and light-off periods (black line) shows significant differences in high-frequency activity (100-250 Hz). P < 0.05, false discovery rate; red asterisks. **C,** Dentate spike counts during light-on and light-off periods of optogenetic stimulation. Wilcoxon paired test, * P = 0.017.

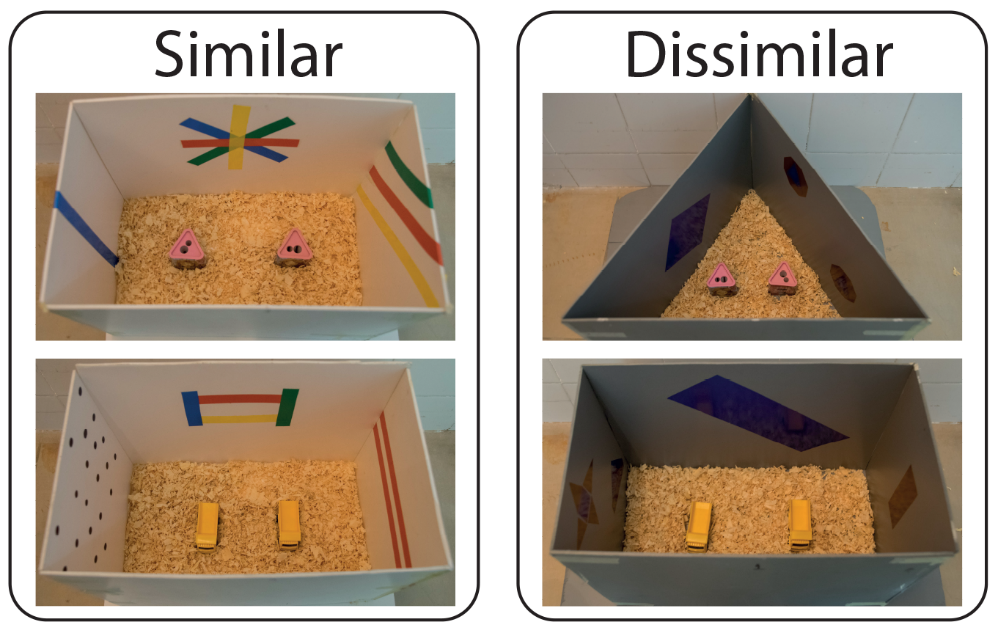

**Supplementary Figure 3. Arenas and objects used for similar and dissimilar object-in-context behavioural tests.** The rectangular arena was 40 cm x 25 cm length x 30 cm high, while the triangular one was 40 cm x 25 cm length x 30 cm high.

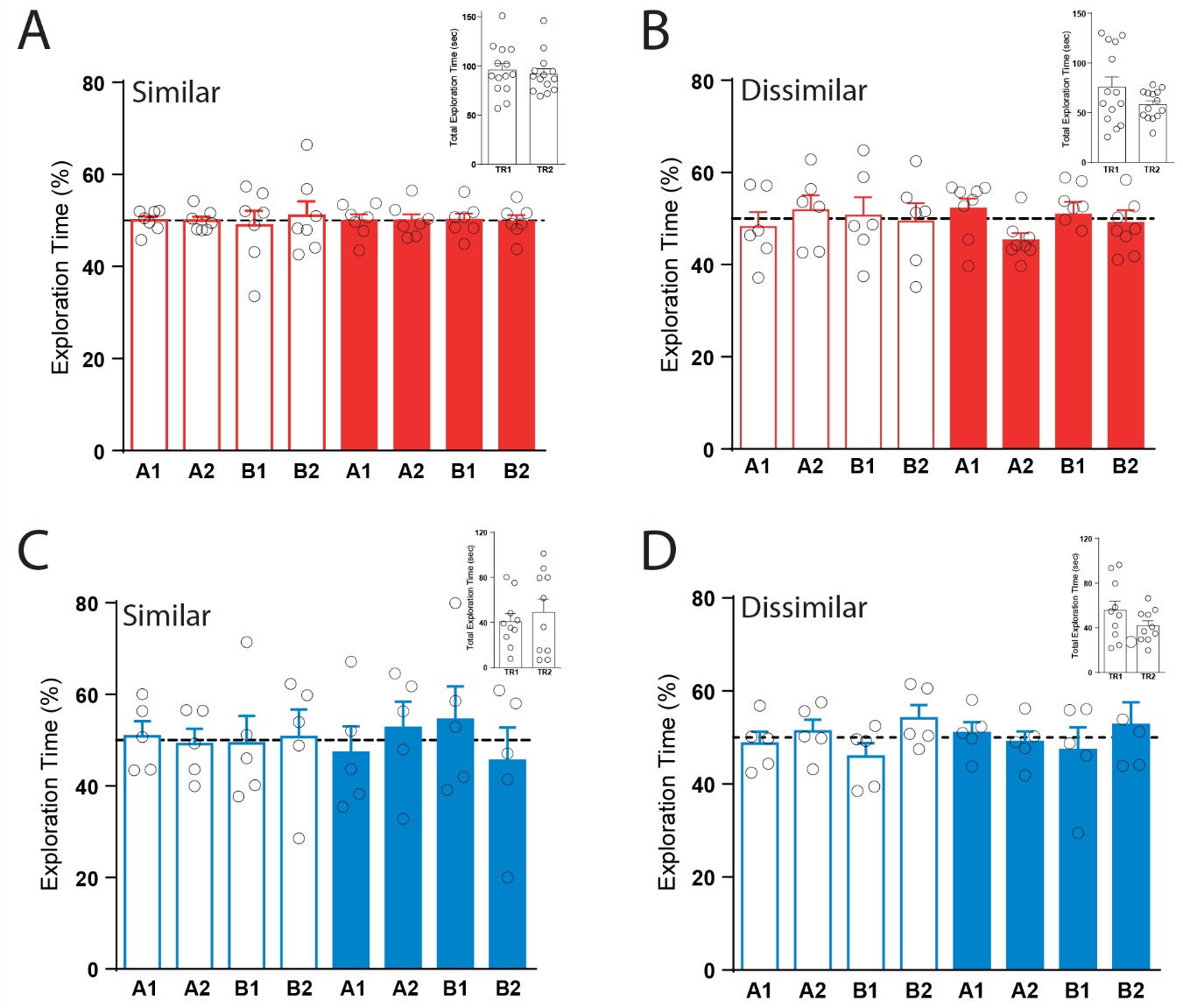

**Supplementary Figure 4. Exploratory behaviour during training sessions in the object-in-context test**. Fraction of time spent in the exploration of each object during training sessions in pharmacological (red) and optogenetic (blue) experiments. Empty bars represent control groups (vehicle, red; NphR-, blue), while solid bars represent experimental groups (DNQX, red; NphR+, blue). **A** and **C** show similar object-in-context tests, and **B** and **D** show dissimilar object-in-context tests. Paired two-way ANOVA; A, B, C, and D, P > 0.05. Insets show the total exploration times of objects during the first (TR1) and the second (TR2) training sessions. Inset: Paired Student´s t test; A, P = 0.45; B, P = 0.16; C, P = 0.49; D, P = 0.18.

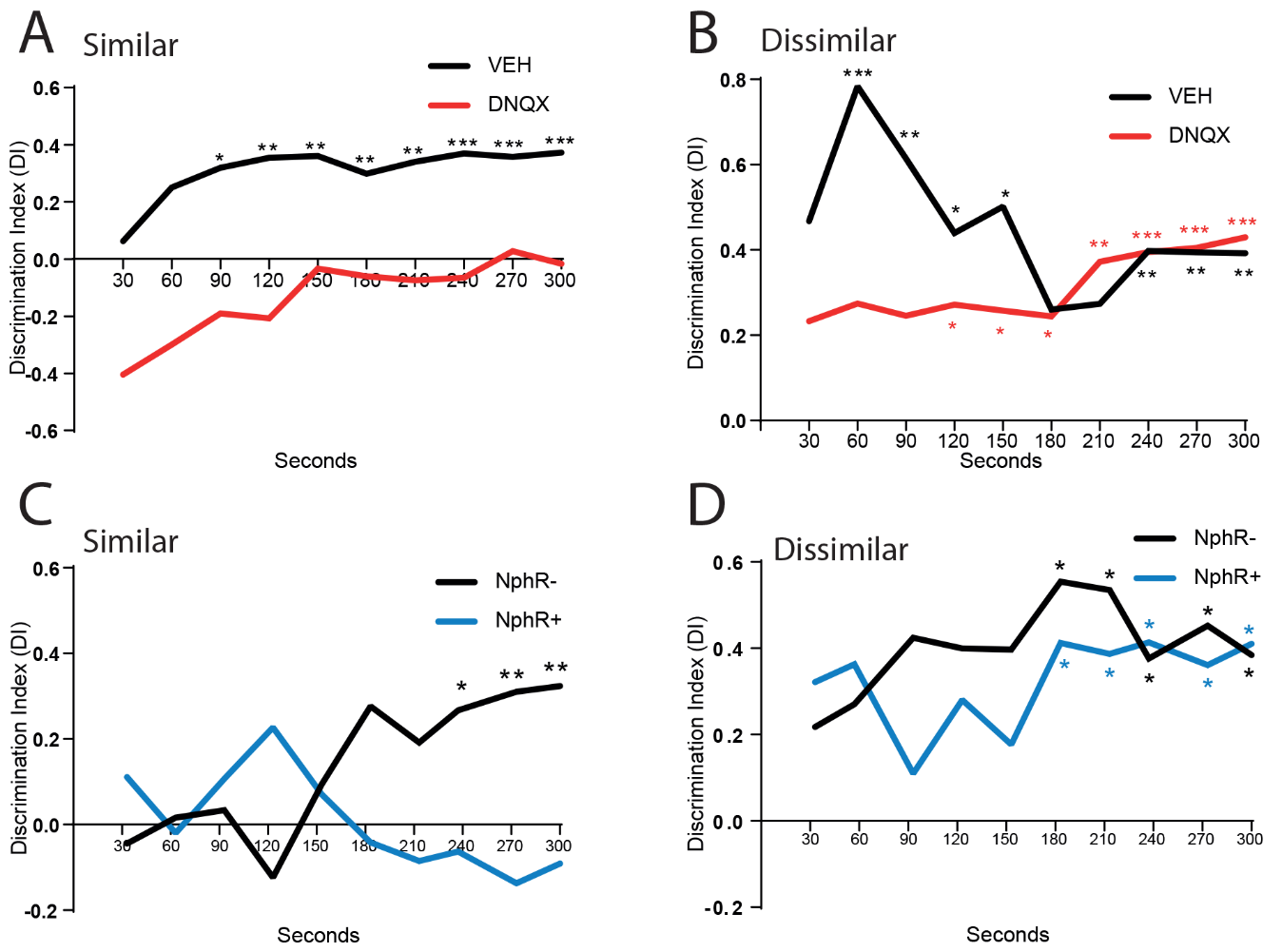

**Supplementary Figure 5. Dynamics of contextual discrimination during the retrieval phase of the object-in-context test.** Black lines represent control groups (vehicle, **A** and **B**; NphR-, **C** and **D**), whereas red and blue lines represent DNQX and NphR+ groups; respectively. Panels **A** and **C** show the similar object-in-context condition, whereas panels **B** and **D** show the dissimilar object-in-context condition. Student´s t test performed point-by-point against zero; *, P < 0.05; **, P < 0.01; ***, P < 0.001.

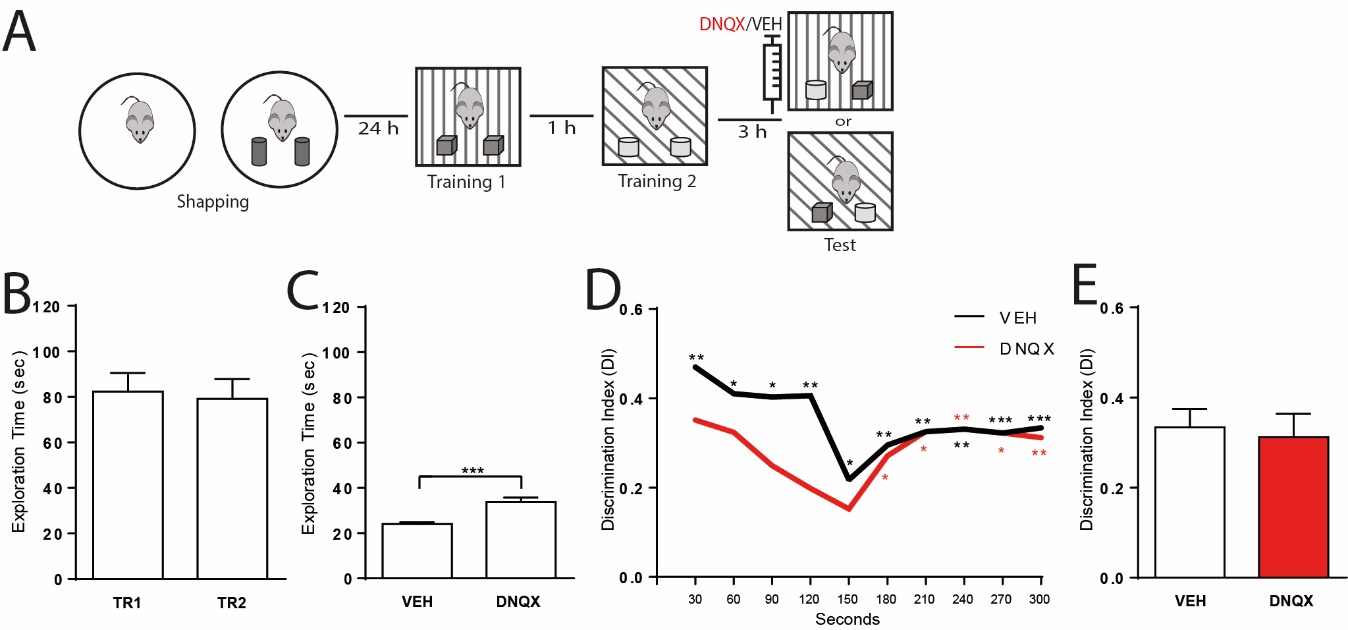

**Supplementary Figure 6. Dentate gyrus AMPA receptors do not regulate memory retrieval in the similar object-in-context behavioural test.** **A,** Schematic representation of drug administration and similar object-in-context behavioural test paradigm. Animals were locally injected in the dentate gyrus with either DNQX or vehicle 15 minutes preceding the retrieval phase. **B,** Total exploration time during the training phase. Student´s t test, P = 0.8. **C,** Total exploration time during the retrieval phase. Student´s t test, ***, P = 0.0008. **D,** Discrimination Index during the retrieval phase for DNQX (red) and vehicle (black) groups. Student´s t test performed point-by-point against zero; *, P < 0.05; **, P < 0.01; ***, P < 0.001. **E,** Cumulative Discrimination Index during the entire retrieval phase. Student´s t test, P = 0.74.

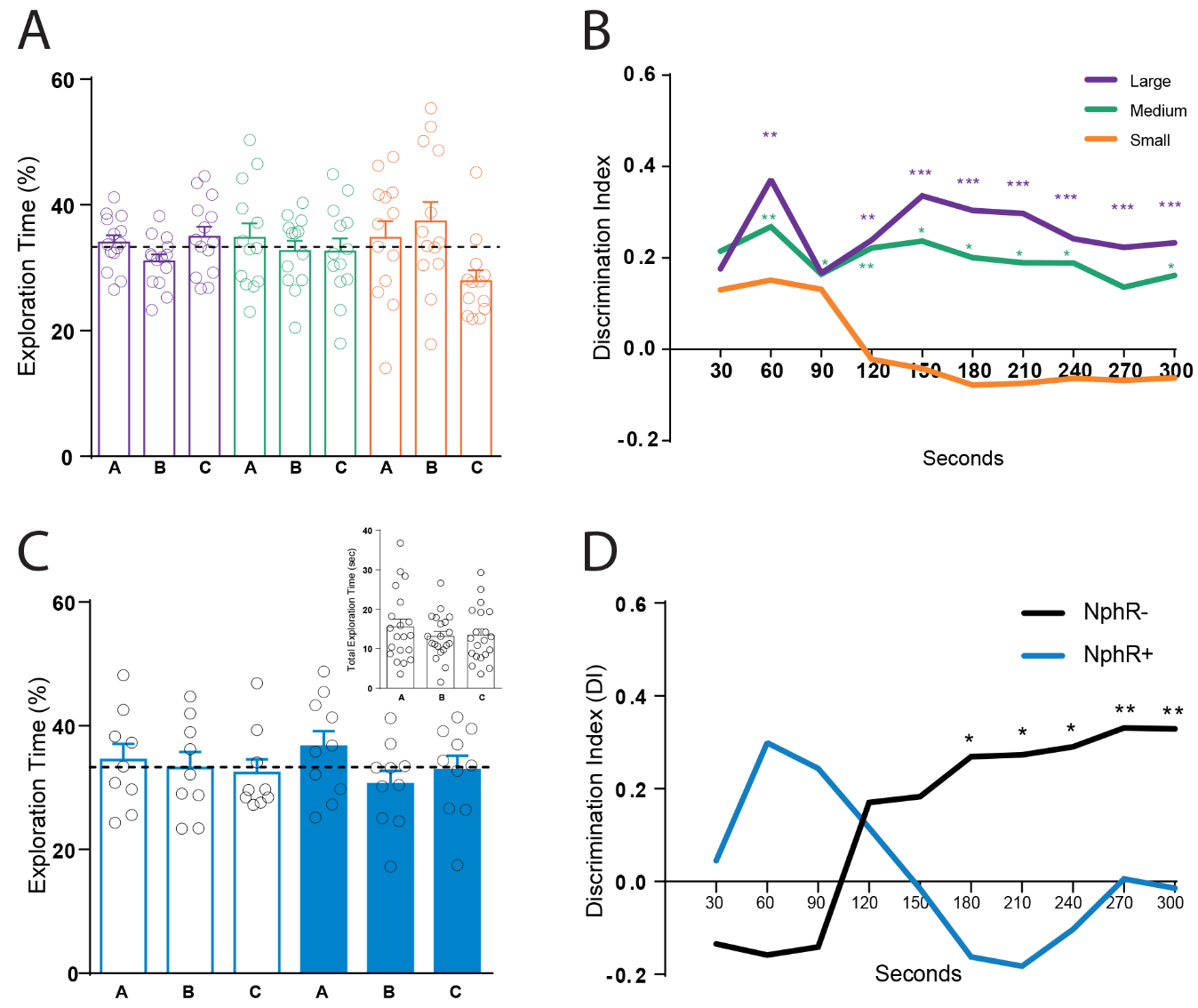

**Supplementary Figure 7. Dynamics of spatial discrimination during the retrieval phase of the spontaneous location recognition test.** **A,** Fraction of time spent in the exploration of each of the three objects (A, B, C) in the different test variations (large, medium, small). Paired two-way ANOVA, P > 0.05. **B,** Dynamics of the Discrimination Index during the retrieval phase in the different variations. Student´s t test performed point-by-point against zero; *, P < 0.05; **, P < 0.01; ***, P < 0.001. **C,** Fraction of time spent in the exploration of each of the three objects (A, B, C) in the different test variations. Empty bars represent NphR- animals, while solid bars represent NphR+ animals. Paired two-way ANOVA, P > 0.05. Inset: total exploration time combining all animals. One-way ANOVA, P = 0.2. **D,** Dynamics of Discrimination Index during the retrieval phase for NphR- and NphR+ groups. Student´s t test performed point-by-point against zero, *, P < 0.05; **, P < 0.01.

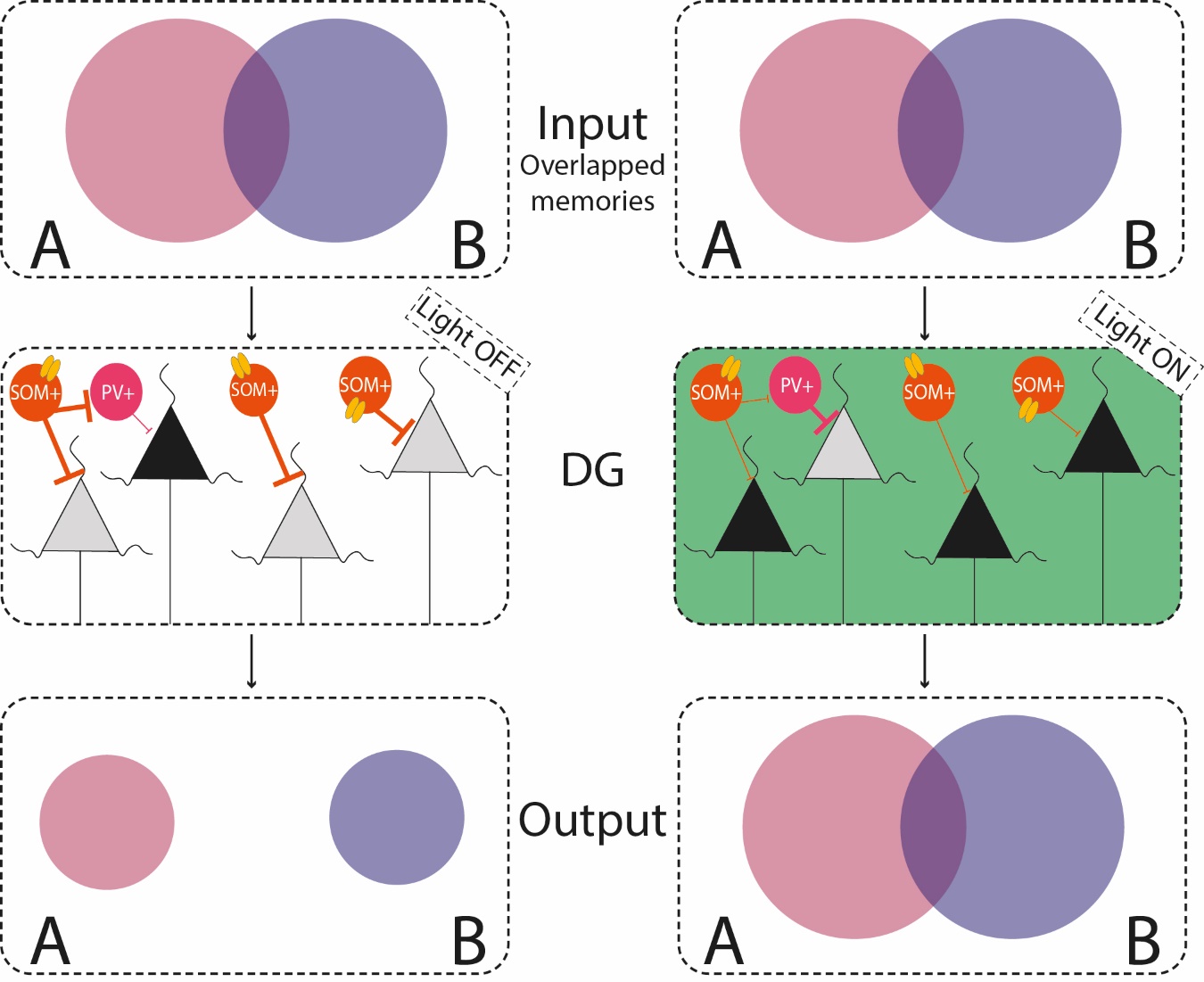

**Supplementary Figure 8. Conceptual model for the role of dentate gyrus somatostatin neurons during pattern separation orthogonalization.** Top, overlapped contextual information recruits the dentate gyrus (DG). Middle, we propose that somatostatin cells (SOM) regulate orthogonalization by directly controlling excitability of granular cells or indirectly by inhibiting parvalbumin cells. Bottom, during successful orthogonalization (left) overlapping input patterns are separated, whereas during SOM suppression by optogenetic stimulation, more granular cells are activated (black cells), thus overlapping inputs are not properly separated.
