## supplementary methods for "Dentate gyrus somatostatin cells are required for contextual discrimination during episodic memory encoding"

Mice were housed at 12 h light/dark cycle at 23°C with food and water ad libitum. Experiments took place during the light phase of the cycle (lights on at 8 AM) in a quiet room located inside the Animal facility with dim light. The experimental protocol for this study was approved by the National Animal Care and Use Committee of the Catholic University of Chile and Favaloro University of Argentina. All experiments were performed on adult (8–30 weeks old) mice.

**Ethic Statement**

All experimental procedures were in accordance with institutional regulations of Institutional Animal Care and Use Committee of the Favaloro University, ASP # 49527/15, Argentinian government regulations (SENASAARS617.2002) and in accordance with Comite Etico Cientifico para el Cuidado de Animales y Ambiente (ID 151223006 and CEBA 13-014) of the Pontificia Universidad Catolica de Chile. All efforts were made to minimize the number of animals used and their suffering.

**Experimental Animals**

For the electrophysiological and optogenetics experiments, three strains of mice were used, C57Bl/6, Ai39(RCL-eNpHR3.0/EYFP) and Sst-IRES-Cre. All transgenic lines were obtained from Jackson laboratories (www.jax.org). We used these strains as controls and refer to them as NpHR− animals throughout the text. Double transgenic animals were obtained from the breeding of SST-IRES-Cre+/+ and Ai39+/- mice, so that they expressed functional Natronomonas pharaonis halorhodopsin (NpHR) exclusively in somatostatin cells. We refer to such animals as NpHR+ throughout the text. For the pharmacological experiments, we use wild-type C57Bl/6 mice from the Pharmacy and Biochemistry School, University of Buenos Aires, Argentina. A total of 95 mice (70 male, 25 female) were used for experiments. We pooled data between female and male mice as discrimination was not significantly different (Unpaired Student´s t test, t = 0.9290, P = 0.3712).

**In vivo electrophysiological recordings under anesthesia**

Anesthesia was induced with isoflurane 4% followed by an intraperitoneal injection of urethane (0.8 g/kg). Animals were left to rest for 20 minutes and placed in a homeothermic blanket that maintained the body temperature at 37° throughout the experiment. After 20 minutes, an intraperitoneal injection of ketamine/xylazine (40/4 mg/kg) was applied. When mice exhibited no reflexes were put into a stereotaxic frame. The skin above the skull was surgically removed and a small craniotomy (1 mm) was drilled above the cortex (coordinates 1.5 mm lateromedial, 2.0 anteroposterior (*1*)). In addition, a customized support bar was glued to the skull to release pressure from the ears and mouth, and hold the animal’s head in position for recordings. At this time, an intraperitoneal cannula was placed to deliver supplementary doses of urethane each 20 minutes (1/12 of the initial volume). After removing the dura over the cortex, the electrode array was placed on the cortex and carefully descended to the dentate gyrus. The electrode array consisted of a matrix of 32 microelectrodes, 50 um apart. An optical fiber (0.1 mm diameter) was associated with the electrodes. The tip of the fiber was 100 um above the most superficial electrode.

**Surgery for chronic implantation and optogenetic stimulation**

Mice were anaesthetized with isoflurane (4% induction, 1.5–2% maintenance) and placed on a stereotaxic frame. Temperature was kept at 37° throughout the procedure (1–2 h) using a homeothermic blanket. The skin was incised to expose the skull. Three or two craniotomies were made with a dental drill for anchoring screws. Additionally, another craniotomy (~1mm) was made above the dentate gyrus bilaterally (anteroposterior -2 mm, mediolateral ± 1.5mm from Bregma). Two optic fibers (diameter 200 um) glued to ceramic ferrules (diameter 230 um) were descended through both craniotomies until reaching the dentate gyrus and fixed in position using dental cement. After surgery, mice received a daily dose of enrofloxacin for 3 days and supplementary analgesia with ketoprofen for 3 days. Animals were allowed 5 days to recover before behavioral training.

**Surgery and drug infusions for pharmacological experiments**

For pharmacological experiments, mice were deeply anesthetized with ketamine/xylazine (150/6.6 mg/kg) and placed in a stereotaxic frame. The skin was incised to expose the skull. Small craneotomy holes were then drilled and a set of 23 G guide cannulae (0.5 cm long) were implanted bilaterally over the dentate gyrus (anteroposterior -2 mm, mediolateral ±1.5 mm from bregma). Cannulae were fixed to the skull with dental acrylic. At the end of surgery, animals were injected with a single dose of meloxicam (0.33 mg/kg) as analgesic and gentamicin (5 mg/kg) as antibiotic. Behavioral procedures were started 5-7 days after surgery. Infusions were made using a 30 G injection cannula connected to a 10 ul Hamilton syringe. Infusions were made on the training day or test day. Cannulated mice received bilateral 0.3 ul infusions of DNQX or vehicle into the dentate gyrus 15 minutes before each training or test session. DNQX was diluted in physiologic solution to a final concentration of 1.89 ug/ul.

**Optogenetic stimulation**

For chronic implants, optogenetic stimulation of dentate gyrus somatostatin interneurons was achieved with a 200 um optic fiber (N.A. 0.37) coupled to a green laser (532 nm, Laserglow Techonologies) that provided a total light power of 0.1–60 mW at the fiber tip. Light stimuli consisted of 5 s light pulses each 15 s, and power at the tip of the fiber was set between 5-15 mW.

For acute recordings, optogenetic stimulation was achieved with an optrode , which consisted of an optic fiber (100 um, N.A. 0.22) attached to an array of electrodes, so electrical recordings and optical stimulation could be performed simultaneously on the same site. Light stimuli consisted of 5 s light pulses each 20 s and power at the tip of the fiber was set between in 5-12 mW.

**Unit crosscorrelation analysis**

Neural activity of dentate gyrus was crosscorrelated with the light pulse. A time window of ± 15 s was defined with point 0 assigned to the light onset. The timestamps of the spikes within the time window were considered as a template and were represented by a vector of spikes relative to t = 0 s, with a time bin of 200 ms and normalized to the basal firing rate of hippocampus neurons. Thus, the central bin of the vector contained the ratio between the number of neural spikes elicited between ± 100 ms and the total number of spikes within the template. Next, the window was shifted to successive light pulses throughout the recording session, and an array of templates was obtained. Then, we calculated the z-score of this crosscorrelogram using the mean and standard deviation obtained, bin-to-bin, from the distribution of 1000 shuffled crosscorrelograms. We classified as excited units, those that presents more than 4 bins with Z-score larger than 3 during laser presentation. Similarly, we classified as inhibited units, those that presents more than 4 bins with z-score more negative than -3 during laser presentation.

**Identification of putative neuron types**

We defined laser-inhibited units as somatostatin cells. To identify different types of units within excited cells we adapted a previous analysis (*2*). Specifically, the trough-to-peak latency and burst index allows to distinguish between glutamatergic cells and GABAergic interneurons, as glutamatergic neurons have longer trough-to-peak latency and higher burst index than GABAergic interneurons. We performed a similar analysis for units that were responsive to laser stiulation. To measure the trough-to-peak, we calculated, for the mean waveform of each detected unit, the temporal difference between the minimum voltage and the maximum voltage (between the minimum and the end of the waveform). To calculate the burst index, first, we calculated the autocorrelogram of the timestamps of each unit outside laser presentation (i.e.; baseline activity); second, we computed the ratio between the peak of the autocorrelogram in the central bins (-1 to 6 ms) and the mean value of the autocorrelogram (200-300 ms); finally, we calculated the logarithm of such ratio. This allowed us to construct a bidimensional vector for excited units. Then, we used hierarchical cluster analysis to discriminate different type of units. The hierarchical cluster analysis showed three types of cluster, two of them were merged in a new cluster because both had similar properties when were compared with somatostatin interneurons. Finally, we compared the trough-to-peak latency and burst index among somatostatin and two clusters were obtained.

**Detection of dentate spikes**

Based on LFP activity, we identified the electrodes located in the hilus and filtered activity between 100-249 Hz. Then, we calculated the z-score of the signal using the mean and the standard deviation of the entire LFP recording. Finally, we selected high-frequency events based on amplitude, with 5-7 SD threshold.

**Arenas and objects used in behavioral tests**

Identical copies of objects made from plastic, glass, or aluminum were used. The height of objects ranged from 4 to 6 cm. All objects were affixed to the floor of the apparatus with an odorless reusable adhesive to prevent them for being displaced during exploration. Objects had no natural relevance for mice as they were not associated with reinforcement. The objects, floor and walls were cleaned with ethanol 10% between sessions. Since no differences were observed in behavioral performance during the experiments between sexes, we pooled animals depending on the genotype or treatment received.

*Object-in-context task*. Four different contexts were used for these experiments. In the dissimilar condition a rectangular and triangular arena were used. Both had homogenous gray walls constructed from opaque foam board. The rectangular apparatus was 40 cm x 25 cm length x 30 cm high, while the triangular one was 40 cm x 25 cm length x 30 cm high. Both contexts had the same surface area to avoid differences due to the size of the arena. Contextual cues were geometric shapes of different colors. In the similar condition, we used two rectangular arenas made of white opaque foam board. The measures of these arenas were identical to those used for the rectangular arena of the dissimilar condition. Then, the cues presented in both contexts were different in shape but same sizes and colors.

*Spontaneous location recognition task.* A circular maze of 40 cm diameter with walls 40 cm high was used. Both floor and wall were gray. Three spatial clues were glued at 15 cm over the floor. A video camera and laser cable were mounted above the maze.

**Object-in-context task**

This behavioral test is composed of three phases that allows the evaluation of the congruency between pairs of context and object (*3*). During the training phase, animals were exposed to two different object-context associations. These two training sessions were 1 hour apart. The test session was carried out twenty-four hours after the training 2. During this phase, a new copy of each of the objects used during the training phase was presented in one of the arenas. The context to be used during the test phase was randomly selected, preserving similar total proportions. Thus, one of the objects was presented in contextual miss-match, the incongruent object, whereas the other object was not, the congruent object. In this task, novelty arises from the novel combination of object and context. Then, exploratory behavior should be driven by retrieval of a particular conjunctive representation of object and context (*4*, *5*).

*Habituation sessions.* These sessions were conducted to familiarize animals with the procedure of being exposed to an environment where they could encounter novel objects. On the first day mice were handled and placed in a circular context and allowed to explore for 10 minutes. Thirty minutes later, they were reintroduced in the arena, yet in this case two different objects were placed in the arena. Mice were allowed to freely explore for 5 minutes.

*Training sessions.* On the first training session, two identical objects (A1 and A2) were placed into one of the arenas (context 1). Animals were placed into the arena and allowed to freely explore the environment for 10 minutes. At the end of the session, animals were returned to their home cage. After 1 hour, animals were exposed to a second object-context association, different from the first one. For that, mice were placed in a different context (context 2), in which a second pair of objects was present (B1 and B2). Arenas were pseudo-randomly assigned as context 1 or 2.

*Test session.* During this session animals were re-exposed to previously familiar context-object pairs for 5 minutes. Mice were reintroduced to context 1 or context 2 where they could explore one copy of object A and one copy of object B. Then depending of the context in which this phase takes place, one of the objects presented will be contextually congruent while the other will be contextually incongruent.

**Spatial Location recognition text**

This behavioral paradigm, as the object-in-context, was comprised by three phases that allowed for the evaluation of spatial location novelty detection (*6*, *7*). During training sessions animals were exposed to three identical objects placed into a circular arena and were allowed to explore them. The separation angle between two of those objects was manipulated in order to generate conditions with variable levels of spatial-location similarity. During the test session, one of the objects was placed in a familiar location, while another copy of the same object was placed in a new location at the middle point between the two previous objects’ locations. The rationale behind the task was that if mice were able to discriminate the two similar spatial locations, their representations should be distinct and resilient to confusion. Thus, mice should show preference to explore the object presented in the novel position during the retrieval phase. However, if the representations of the two similar locations were not sufficiently segregated, then mice should behave as if the new location was familiar.

*Habituation sessions.* These sessions were conducted to familiarize animals with the procedure of being exposed to an environment where they could encounter novel objects. During these sessions, animals had the opportunity to generate a spatial map of the environment. To that end, mice were repeatedly exposed to the environment for 10 minutes during four consecutive days.

*Training session.* During this session, animals were placed in the circular arena where three identical objects had been previously positioned. The angle between two of the objects was changed depending on the variation in use (large, 180 degrees; medium, 120 degrees; or small, 50 degrees). Mice were then allowed to explore the environment for 10 minutes.

*Test session.* In this session only 2 copies of the previously presented objects were placed in the arena. One of the objects was located in the same position as before, yet the other object was placed in the intermediate position occupied by the two objects in the previous session. This session lasted 5 minutes.

**Behavioral Analysis**

For each behavioural session we quantified the exploration time of each object. For the test phase, we analyzed the exploration time for every copy of the object using a Matlab plug-in (ID tracker). For the training phase, manual score was performed. For the object-in-context test session we calculated the Discrimination Index (DI) as t_incongruent_ – t_congruent_ / t_total exploration_ of the session. For the spontaneous location recognition test, the DI was calculated as t_novel-position_– t_familiar-position_ / t_total exploration_. For all experiments, object exploration was defined as the mouse having its nose directed to the object and located at a distance of 2 cm or less. Climbing over or sitting on the objects was not considered as exploration. Two persons scored independently the videos; one of them was blind to all conditions.

**Statistical analysis**

Statistical analyses were performed with GraphPad 6.01 and Matlab. We used parametric analysis depending when data distributed normally. Electrophysiological data were analyzed using Kruskal-Wallis followed by Tukey-Kramer multiple comparison test. Behavioral data were analyzed using two-tailed Student’s t test or two-tailed Wilcoxon test. For comparisons between two repeated-measured groups two-tail paired Student's T test or two-tailed paired Wilcoxon test were used. For more than three groups, we performed One-way ANOVA followed by Tukey's post-hoc test or Kruskal-Wallis test followed by Dunn's post-hoc test. Two-way ANOVA followed by Tukey's post-test was used when three or more groups were involved. In all cases, P-values were considered statistically significant when smaller than 0.05. All data are presented as the mean ± s.e.m.

**Histology and Immunocytochemistry**

At the end of electrophysiological recordings and behavioural testing, mice were terminally anesthetized and intracardially perfused with saline followed by 20-min fixation with 4% paraformaldehyde. Brains were extracted and post-fixed in paraformaldehyde for a minimum of 24 h before being transferred to PBS azide and sectioned coronally (70–100 um thickness). Sections were further Nissl-stained. Location of electrode shanks and optical fibers were determined in reference to standard brain atlas coordinates (*1*) under a light transmission microscope.

**Spike Sorting**

Semiautomatic clustering was performed by KlustaKwik (*8*). This method was applied over the 32 channels of the silicon probe, grouped in 8 pseudo-tetrodes of 4 nearby channels.
